# Endothelial-specific Gata3 expression is required for haematopoietic stem cell generation

**DOI:** 10.1101/2021.09.14.460344

**Authors:** Nada Zaidan, Evangelia Diamanti, Leslie Nitsche, Antonella Fidanza, Nicola K. Wilson, Lesley M. Forrester, Berthold Göttgens, Katrin Ottersbach

**Affiliations:** Centre for Regenerative Medicine, Institute for Regeneration and Repair, University of Edinburgh, Edinburgh, EH16 4UU, UK; Department of Haematology, Wellcome Trust-Medical Research Council Cambridge Stem Cell Institute, University of Cambridge, Cambridge CB2 0AW, UK

**Keywords:** haematopoietic stem cell, Gata3, haemogenic endothelial cells, endothelial-to-haematopoietic transition

## Abstract

To generate sufficient numbers of transplantable haematopoietic stem cells (HSCs) *in vitro*, a detailed understanding of how this process takes place *in vivo* is essential. The endothelial-to-haematopoietic transition (EHT), which culminates in the production of the first HSCs, is a highly complex process during which key regulators are switched on and off at precise moments and which is embedded into a myriad of microenvironmental signals from surrounding cells and tissues. We have previously demonstrated an HSC-supportive function for Gata3 within the sympathetic nervous system and the sub-aortic mesenchyme, but show here that it also plays a cell-intrinsic role during the EHT. It is expressed in haemogenic endothelial cells and early HSC precursors, where its expression correlates with a more quiescent state. Importantly, endothelial-specific deletion of Gata3 shows that it is functionally required for these cells to mature into HSCs, placing Gata3 at the core of the EHT regulatory network.

## Introduction

Due to its central position in the generation of haematopoietic stem cells (HSCs), the endothelial-to-haematopoietic transition (EHT) has been studied in considerable detail, using a number of different systems and species, such as chicken, mouse, human, zebrafish and pluripotent stem cells (PSCs). Particularly the latter has been the focus of many labs as the ultimate goal is to be able to generate HSCs *in vitro* for clinical applications. Despite recent successes ^1^, this remains challenging and further dissection of the EHT as it takes place *in vivo* is required.

The generation of the first transplantable HSCs initiates in the dorsal aorta of the aorta-gonad-mesonephors (AGM) region at E10.5 in mouse embryos ^2, 3, 4^. It involves activating a haematopoietic transcriptional program in a subset of endothelial cells (ECs), as a result of a number of internal and external signals, which drives these cells to adopting a haematopoietic fate (recently reviewed in ^5^). These so-called haemogenic endothelial cells (HECs) undergo major morphological changes that allow them to break tight junctions to neighbouring ECs, adopt a haematopoietic fate and translocate to the lumen of the aorta where they will eventually be released into the circulation and go on to colonise the foetal liver. This process results in the appearance of intra-aortic cell clusters in which HECs undergo further maturation steps from pro-HSCs to type I and type II pre-HSCs, before becoming fully mature HSCs ^6^. While it is known that the two transcription factors Runx1 and Gata2 are essential for the EHT to take place, without which HSCs fail to be generated ^7, 8^, it is clear that they do not act in isolation. For example, Sox17 is required upstream for arterial fate and HEC specification ^9, 10^, while Gfi1 and Gfi1b are downstream targets of Runx1, necessary for EHT completion ^11^. It has even been suggested that the relative levels of Sox17 and Runx1 determine the fate of an EC, with a higher Runx1:Sox17 tipping the balance towards the haematopoietic fate ^12^; however, how all of these transcription factors interact and whether they form multi-component complexes remains unknown.

A number of single-cell RNA sequencing datasets have been generated from mouse and human embryos ^13, 14, 15, 16, 17, 18^, which have revealed further intermediate endothelial and HSC precursor populations and have provided some insight into the regulation of the EHT. These studies have identified new markers for HECs, such as CD44 ^16, 17^ and Neurl3 ^15^, providing improved purification strategies, and have demonstrated a functional role for the mTOR signalling pathway ^18^, CD44-hyaluronan interaction ^16^ and endothelin signalling ^14^ in the EHT.

Many other details of the EHT, however, remain to be discovered, including why only some of the aortic ECs become haemogenic, what the mechanical forces and structural proteins are that bring about the morphological changes, how this is driven metabolically and, possibly, by cell cycle changes, and the role of the local microenvironment.

In this study, we reveal the transcription factor Gata3 as another important regulator of the EHT. Its expression is upregulated in HECs and early HSC precursors, thereby enriching for haemogenic potential, but is downregulated before fully mature HSCs are formed. Through RNA-Seq analysis we show co-expression of Gata3 with many important EHT regulators and have been able to link Gata3 expression with a more quiescent cell state. Importantly, endothelial-specific deletion of Gata3 significantly reduces haematopoietic stem and progenitor (HSPC) formation, which, together with its haematopoiesis-supportive role in the co-developing sympathetic nervous system (SNS) ^19^ and the sub-aortic mesenchyme ^20^, gives Gata3 a multi-faceted, central role in HSC generation.

## Results

### Gata3 is expressed in a subset of endothelial and haematopoietic cells

We had previously detected Gata3 expression in a subset of ECs ^19^ (**Figure 1A, red arrowheads**), often in the vicinity of intra-aortic cell clusters (**Figure 1A, yellow arrowheads**), which suggested that Gata3 may be expressed in HECs. We decided to carry out a more careful analysis using a mouse line that carries a GFP reporter inserted into the *Gata3* locus ^21^. Immunohistochemical co-staining for Gata3 with an anti-Gata3 antibody and an anti-GFP antibody confirmed this construct to be a faithful reporter of endogenous Gata3 expression (**Figure S1**), that recapitulated the expression of Gata3 within individual endothelial cells (**Figure 1B, arrows**) and at the base of intra-aortic clusters (**Figure 1B, yellow arrowhead**), but not within clusters. Flow cytometry analysis confirmed that a fraction (6.4%) of ECs (VE-Cadherin [VEC]+) expresses Gata3-GFP at E10.5 when HEC frequency is at its highest (**Figure 1C**). Interestingly, more than 40% of these Gata3-GFP+ ECs also express the hematopoietic markers CD45 (late haematopoietic differentiation marker) and/or CD41 (early marker), with 19% expressing both. The percentage of CD45+ Gata3-GFP+ cells was much lower within the VEC-fraction, suggesting that the majority of Gata3-GFP+ cells that express haematopoietic markers are early VEC+ hematopoietic stem and progenitor cells (HSPCs). The percentage of Gata3-expressing CD45+ cells (both within the VEC+ and VEC-fraction) increases substantially at E11.5 (**Figure 1D**).

**Figure 1.**
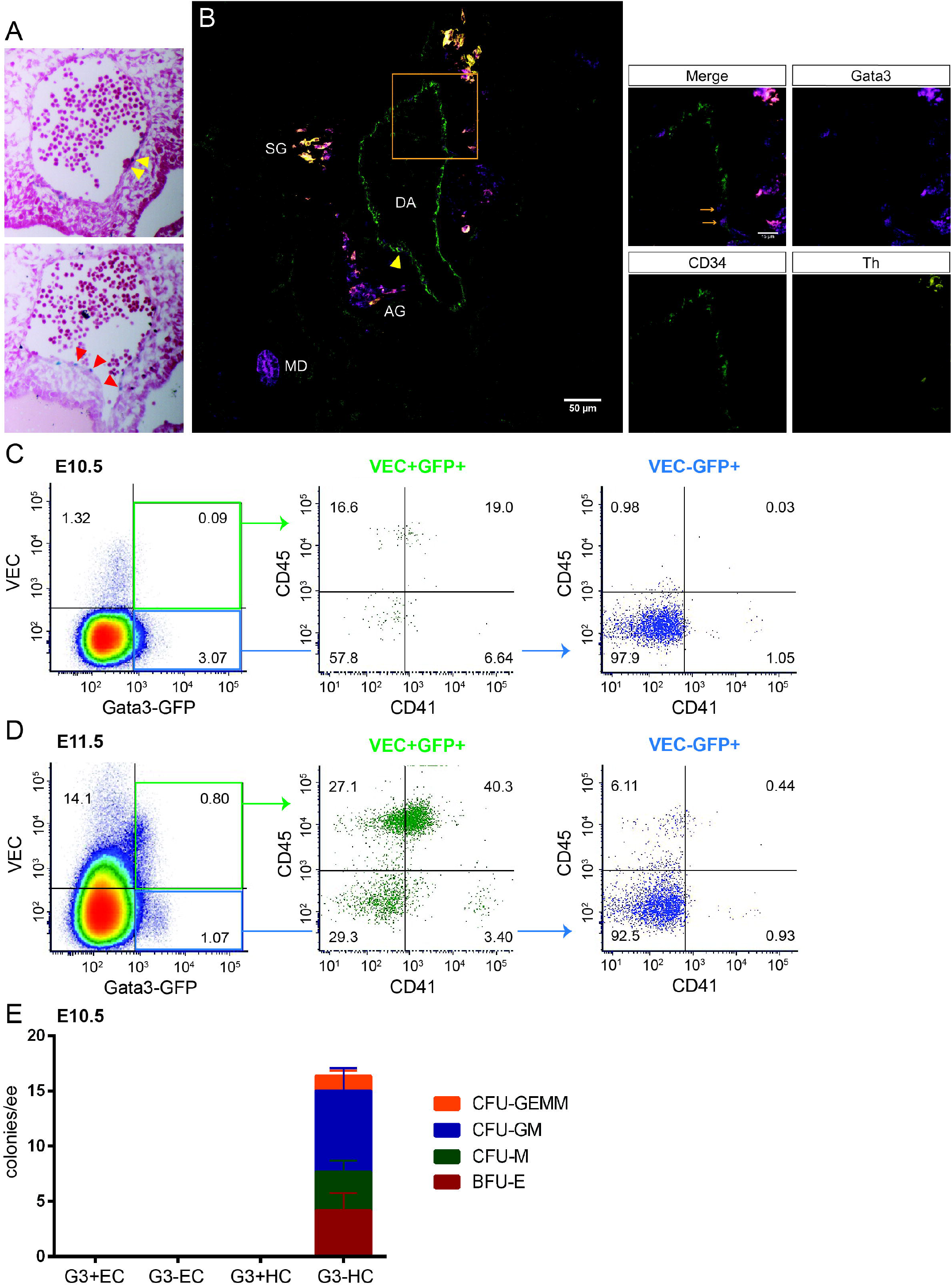
Gata3 is expressed in a subset of endothelial and haematopoietic cells. (**A**) Cryosections of E10.5 *Gata3*^*lz/+*^ with Gata3-LacZ expression shown in blue through Xgal and counterstained with Neutral Red. Yellow arrowheads point to Gata3+ cells near intra-aortic clusters and red arrowheads highlight Gata3+ endothelial cells. (**B**) Section from a Gata3-GFP+ E11.5 embryo stained with CD34 (green), Gata3-GFP (magenta) and Th (yellow). Orange box indicates area shown at higher magnification on the right. Orange arrows highlight Gata3+ cells inside the endothelial layer. AG: adrenal anlage, DA: dorsal aorta, SG: sympathetic ganglia, MD: mesonephric duct. Flow cytometry analysis of G3-GFP+ samples from E10.5 (**C**) and E11.5 (**D**) AGMs, stained for GFP, Cdh5, CD41 and CD45. (**E**) CFU-C assay representing the progenitor counts of freshly sorted and directly plated G3-EC: CD41/45-GFP-VEC+, G3+EC: CD41/45-GFP+VEC+, G3-HC: CD41/45+GFP-, and G3+HC: CD41/45+GFP+.

To determine if the Gata3-GFP+ haematopoietic population at E10.5 does indeed contain early progenitors, we plated them directly in colony-forming (CFU-C) assays alongside Gata3-GFP-haematopoietic cells (HCs) and Gata3-GFP+/-ECs. As expected, ECs do not produce haematopoietic colonies when plated directly in methylcellulose (**Figure 1E**). Interestingly, only the Gata3-GFP-haematopoietic population gave rise to colonies, but not the GFP+ fraction, suggesting that the latter consists either of mature haematopoietic cells or of very early precursors that require further maturation towards the haematopoietic fate.

### Gata3 expression enriches for haemogenic endothelial activity

Inherent haemogenic potential can be revealed via a co-culture step on OP9 stromal cells ^22^. To test whether Gata3 expression marks HECs, Gata3-GFP+/-ECs were sorted (**Figure S2A**) and cultured on a layer of OP9 cells. At the end of the culture, cells were assessed for haematopoietic marker expression by flow cytometry and for progenitor potential by CFU-C assay (**Figure 2A**). Both endothelial populations were able to give rise to haematopoietic cells during the co-culture (**Figure 2B**), with a trend towards a higher production of CD45+ cells from Gata3-GFP+ ECs that did not quite reach significance (**Figure 2C**). In functional CFU-C assays following co-culture, however, Gata3-GFP+ ECs had a noticeably higher colony output (**Figure 2D**), which was highly significant for all colony types, including CFU-GEMM, produced by the most immature progenitors. These results demonstrate that Gata3 expression enriches for HECs.

**Figure 2.**
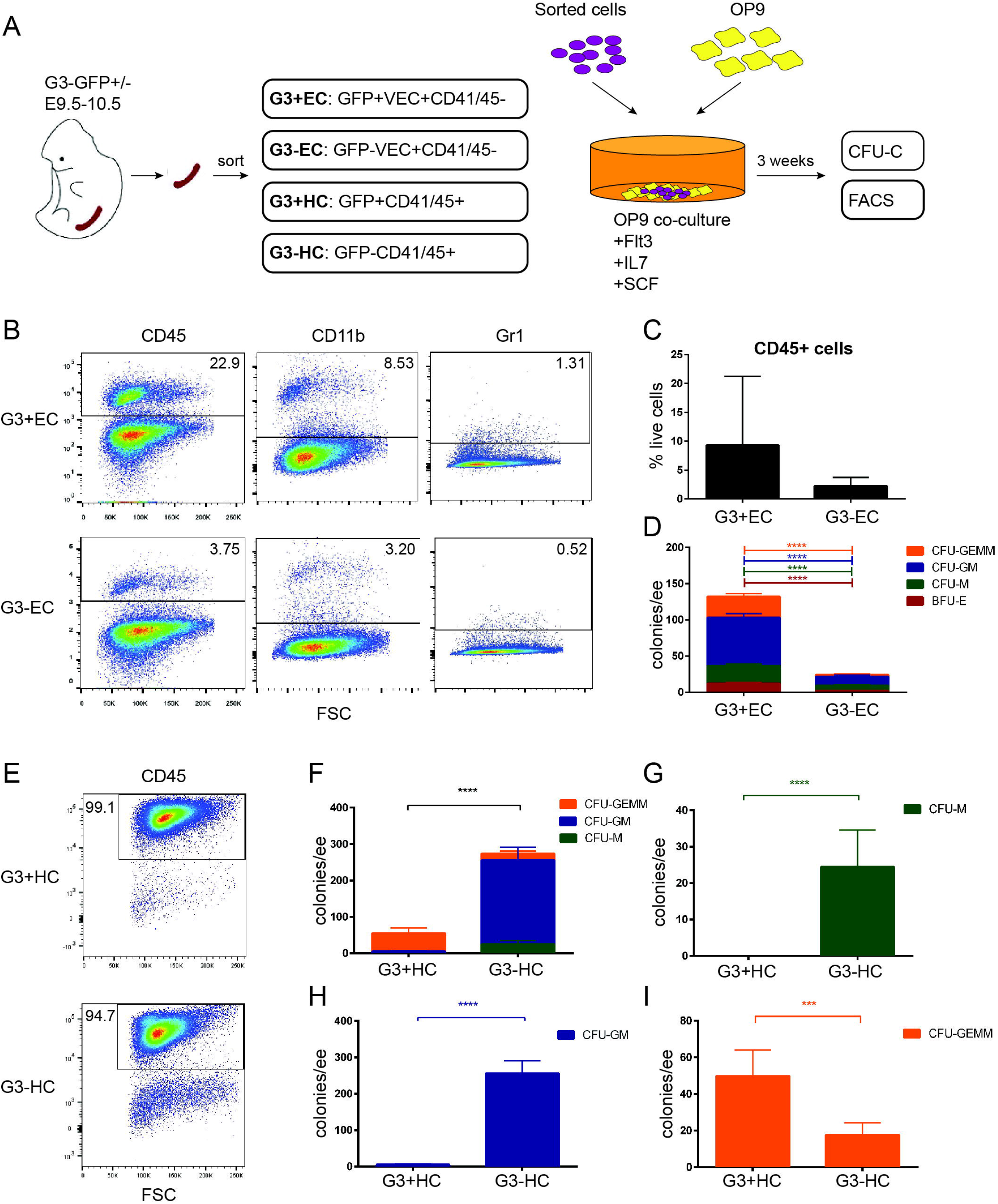
Gata3 enriches for haemogenic endothelial cells. (**A**) Outline of co-culture experiments. (**B**) Flow cytometry analysis of haematopoietic output from Gata3+/-ECs at the end of the co-culture. Total CD45+ cells (as detected by flow cytometry) (**C**) and haematopoietic progenitors (as detected by CFU-C assays) (**D**) produced by Gata3+/-ECs at the end of the co-culture. (**E**) Flow cytometry analysis of CD45+ cells produced from Gata3+/-HCs at the end of the co-culture. Total progenitors (**F**), CFU-M (**G**), CFU-GM (**H**) and CFU-GEMM (**I**) (as detected by CFU-C assays) produced by Gata3+/-HCs at the end of the co-culture. *** p<0.001; **** p<0.0001; Mann-Whitney test.

### Gata3 marks haematopoietic stem cell precursors

We also tested the potential of the Gata3-GFP+/-haematopoietic populations (**Figure 2A, S2A**) in these OP9 co-cultures. As these cells started off as being CD45 and/or CD41-positive, it was not surprising that they gave an almost entirely CD45+ output (**Figure 2E**). Interestingly, while overall progenitor potential was higher in the Gata3-GFP-fraction (**Figure 2F**), splitting this into the individual progenitor types revealed that Gata3 expression enriches specifically for the most immature CFU-GEMM progenitor (**Figure 2G-I**), suggesting that Gata3-GFP may be marking HSC precursors.

The intermediate steps involved in the maturation of HEC into transplantable HSCs have been carefully dissected in recent years with the help of a culture system developed in the Medvinsky lab ^6, 23^. It involves aggregation of isolated cell populations with OP9 cells and culturing these co-aggregates as explants in the presence of cytokines. At the end of the culture, the potential of the initial populations is assessed by flow cytometry analysis and transplantations (**Figure 3A**). Depending on the cell surface marker combination and the embryonic stage, precursor populations are defined as pro-HSCs, pre-HSCs type I and pre-HSCs type II ^6^. Pro-HSCs are restricted to the E9 AGM, while type I pre-HSCs are detected at E10-11. Pre-HSC type II only emerge at E11. To test whether Gata3 is expressed in HSC precursors, we sorted Gata3-GFP +/-ECs and HCs (**Figure S2B**) and cultured them as OP9 co-aggregates (**Figure 3A**). As in the co-culture systems, all four populations were able to generate CD45+ HCs, with the sorted haematopoietic fractions producing substantially more CD45+ cells than the EC populations (**Figure 3B**). Endothelial cells are unable to transdifferentiate into transplantable HSCs in these culture conditions ^6^, and indeed neither Gata3+ nor Gata3-ECs produced any detectable chimerism in transplant recipients (**Figure 3C,D**). Intriguingly, repopulation activity at E9.5-E10.5 was restricted to the Gata3-GFP+ haematopoietic fraction, which completely shifted to the Gata3-GFP-haematopoietic fraction at E11.5. All of the results taken together with our previous data ^19^ imply that Gata3 expression is switched on in HECs, with Gata3 expression continuing during the early stages of HSC maturation, but being switched off at the pre-HSC type II stage and remaining off in emerging AGM HSCs.

**Figure 3.**
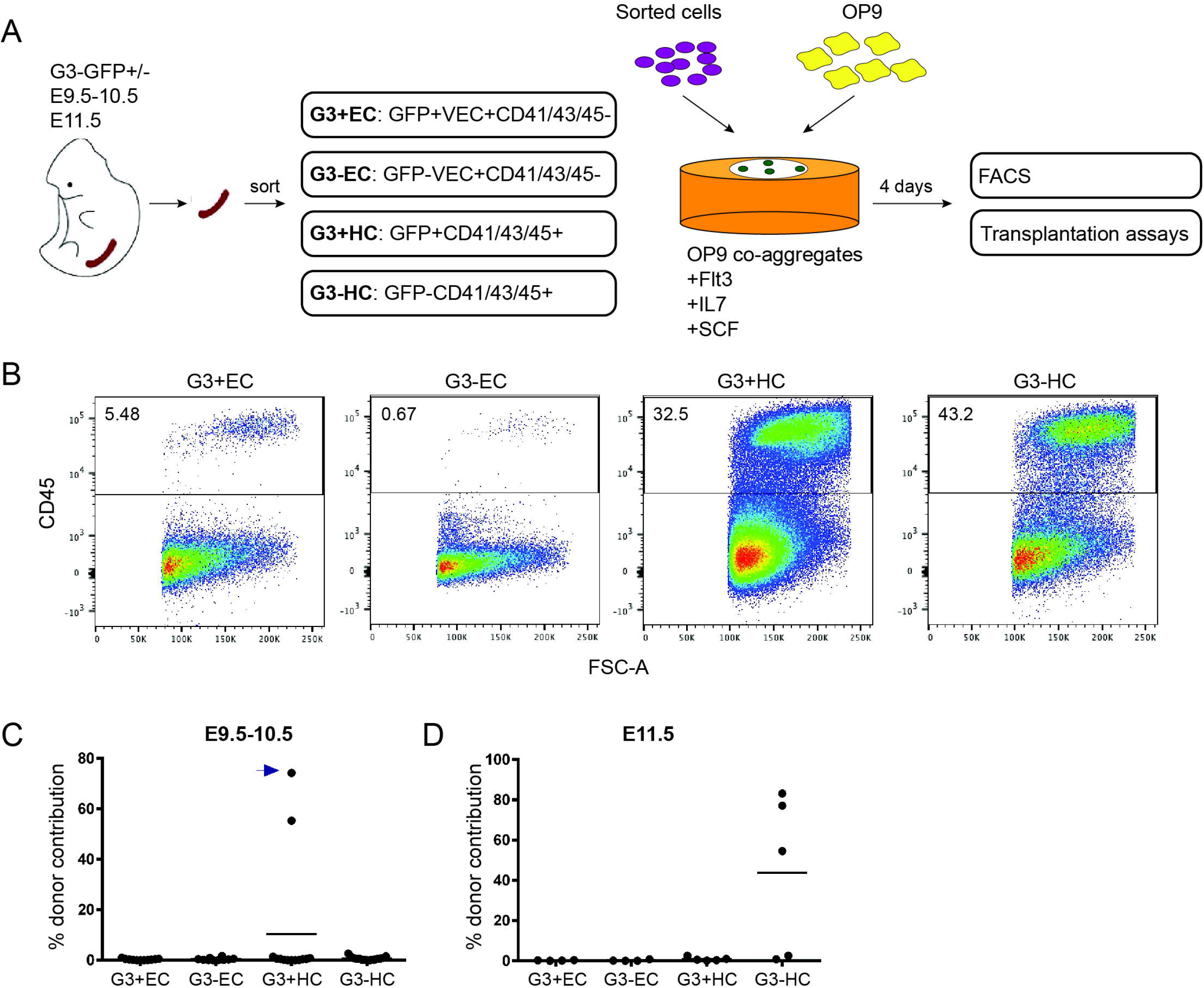
Gata3 is expressed in early HSC precursors. (**A**) Outline of co-aggregate experiments. (**B**) Total CD45+ cell output (as measured by flow cytometry) from E9.5-10.5 Gata3+/-ECs and HCs. Percent Donor contribution after 4 months in recipients of co-aggregates from E9.5-10.5 (**C**) and E11.5 (**D**) Gata3+/-ECs and HCs. Arrow highlights data point from analysis after 1 month as this recipient died unexpectedly before the 4 months analysis.

### Gata3-expressing cells show an enrichment for key regulators of EHT

To get a better understanding of how Gata3 may be involved in the EHT and the maturation of pre-HSCs, we performed RNA-Seq on small pools (20 cells/pool; 20 pools/population) of Gata3-GFP+ ECs and HCs and compared their transcriptome to the Gata3-GFP-fraction from the same cell compartments (**Figure 4A**). Principal Component Analysis (PCA) showed a clear separation of ECs from HCs along component 1, with each cell population displaying a distinct subdivision according to Gata3 expression along component 2 (**Figure 4B**). To gain further insight into the potential role of Gata3 and what defines the Gata3-expressing fraction within each cell compartment, we identified genes that are differentially expressed between Gata3-positive and negative cells. A higher number of genes were upregulated in Gata3-positive cells (EC: 557; HC: 1527) than were downregulated (EC: 253; HC: 232), with some overlap between the two cell types (**Figure 4C; Table S1**). Gene ontology analysis of the upregulated genes saw a significant enrichment of processes associated with stem cell development and differentiation and migration in both cell populations, which may be a reflection of these cells undergoing morphological changes during EHT (**Figure S3**). Genes that were downregulated in the Gata3+ haematopoietic fraction were largely associated with differentiated blood lineages (e.g. *Icos, Irf8, Klf4, Ccl3 and Il6ra*), which reflects the HSC precursor status of the Gata3+ cells and the fact that the more mature HCs will be included in the Gata3-fraction. Interestingly, Gata3+ ECs displayed a downregulation of genes linked to the positive regulation of cell cycle and proliferation.

**Figure 4.**
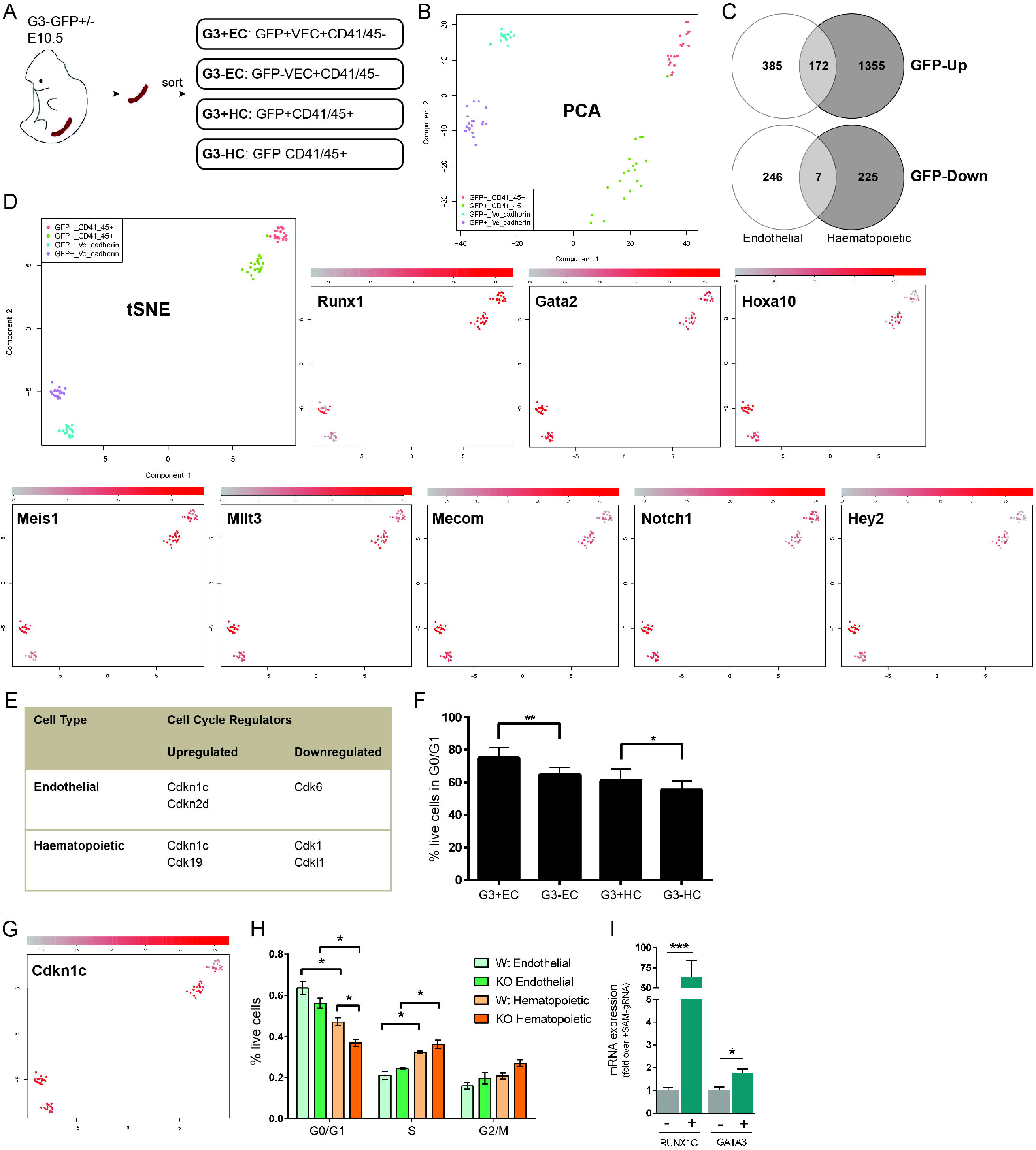
Gata3 expression correlates with other EHT regulators and marks quiescent cells. (**A)** Cell populations sorted and analysed by RNA-Seq. (**B**) Principal Component Analysis (PCA) of individual samples of the 4 cell populations. (**C**) Venn diagrams of genes upregulated and downregulated in Gata-GFP+ ECs and HCs. (**D**) tSNE plots of the 4 cell populations pseudo-coloured for the expression of the indicated genes. (**E**) Cell cycle regulators differentially expressed in Gata3-GFP+ ECs and HCs. (**F**) Percentage of quiescent cells in the Gata3-GFP+/-EC and HC populations. * p<0.05; ** p<0.01; paired t-test. (**G**) tSNE plot pseudo-coloured for the expression of *Cdkn1c* in the 4 cell populations. (**H**) Percentage of *Cdkn1c* wild-type and knockout ECs and HCs in the different phases of the cell cycle. * p<0.05; paired t-test. (**I**) Expression of *RUNX1C* and *GATA3* as measure by qPCR in hiPSCs with (+) and without (-) *RUNX1C* activating specific gRNA as part of the UniSAM endogenous gene activation system. * p<0.05; *** p<0.001; Mann-Whitney test.

To better understand the position of Gata3 in the network of EHT regulators, we analysed the expression of a number of key regulators of HSC generation in our datasets. Both Runx1 and Gata2 are transcription factors known to be essential for the transition of HECs into HSCs in the AGM ^7, 8^. While *Gata2* was uniformly and significantly upregulated in the Gata3+ EC fraction (**Figure 4D, Table S1**), *Runx1* was restricted to a subset of Gata3+ EC (**Figure 4D**). Interestingly, neither of them was differentially expressed between the Gata3+ and Gata3-HC population, reflecting their continued expression throughout HSC maturation, while Gata3 becomes rapidly downregulated before HSCs fully mature.

Among the genes commonly upregulated in Gata3-GFP+ ECs and HCs (**Table S1**) were well-known HSC regulators such as *Hoxa10*, which was recently shown to drive the transition of HECs into HSCs derived from human iPSCs ^1^, *Meis1*, which promotes the reprogramming of mature haematopoietic progenitors to HSCs ^24^, and *Mllt3*, which is essential for human HSC self-renewal ^25^ and was reported to be upregulated in cells undergoing EHT ^16^ (**Figure 4D**). *Mecom*, another transcription factor essential for foetal and adult HSC function ^26^, that was recently, together with *Hoxa10* and *Meis1*, described as a marker for E9.5 aortic haemogenic endothelium ^27^, was upregulated in the Gata3-GFP+ EC fraction. The Notch signalling pathway is essential for the EHT; however, its activity needs to be downregulated again for fully mature HSCs to emerge ^28^, which is reminiscent of the downregulation of *Gata3* after the HEC and early HSC precursor stage shown here. Indeed, both *Notch1* and one of its important downstream targets in this process, *Hey2*, were upregulated in the Gata3-GFP+ EC and HC fraction. This co-regulation of *Gata3* with important drivers of the EHT further confirms its importance in this process.

### Gata3 expression correlates with a more quiescent cell state

Among the most prominent gene ontology terms associated with the genes downregulated in Gata3-GFP+ ECs were proliferation and positive regulation of the cell cycle (**Figure S3**). We interrogated the differentially expressed genes specifically for cell cycle regulators and uncovered as a general trend an upregulation of cell cycle inhibitors and a downregulation of cell cycle promoters in the Gata3-GFP+ fractions (**Figure 4E**). Looking at the cell cycle status of these cells, we also observed a significant enrichment of quiescent cells in the Gata3-GFP+ subsets (**Figure 4F**). Since the cell cycle inhibitor *Cdkn1c* was upregulated in both Gata3-GFP+ cell types (**Figure 4E,G; Table S1**), we hypothesised that it was responsible for the quiescent phenotype. Indeed, there was a decrease in Gata3-GFP+ ECs and HCs in the G0/G1 phase of the cell cycle from Cdkn1c-deficient embryos (**Figure 4H**). We also noticed that Gata3-GFP+ ECs were generally more quiescent than HCs, suggesting that HECs may exit the cell cycle to undergo the massive morphological changes required for their transition into HCs, with emerging HCs then starting to proliferate to expand the pre-HSC pool, as has been observed by others ^16, 29, 30^. Interestingly, an upregulation of RUNX1C in human PSCs has recently been associated with an exit from the cell cycle ^31^. These RUNX1C-expressing cells also had higher levels of GATA3 (**Figure 4I**), which not only confirms the association of Gata3 expression with a more quiescent cell state, but may also point to a direct link between these two EHT regulators.

### Gata3 is required for the endothelial-to-haematopoietic transition

As we had detected Gata3 expression in HEC and HSC precursors, we wanted to confirm whether it plays a functional role during the EHT. We therefore crossed a conditional Gata3 knockout mouse line ^32^ with a VEC-Cre line ^7^ to delete *Gata3* specifically within the endothelial lineage and its derivatives and assessed how this affected HSPC numbers in the AGM (**Figure 5A**). Heterozygous and homozygous deletion of Gata3 significantly reduced progenitor numbers in the AGM at both E10.5 and E11.5, demonstrating that endothelial-specific Gata3 expression is required for haematopoietic progenitor formation and that this is dose dependent (**Figure 5B,C**). A complete knockout of Gata3 resulted in a very similar reduction of progenitors, showing that it is the expression of Gata3 in endothelial cells that is important in this context (**Figure S4A**,**B**). We had previously reported that explant-culturing of AGMs rescued the HSC defect in *Gata3*^*+/-*^ embryos ^19^. To see if that could also rescue the progenitor defect observed here, we inserted a 3-day explant culture step before plating AGM cells in methylcellulose. Explant-culturing AGMs prior to CFU-C assays also gives a more accurate estimate of progenitor numbers generated specifically in the AGM, as a proportion of the progenitors detected in freshly dissected AGMs are likely to be yolk sac derived. Progenitor numbers in VEC-Cre+ *Gata3*^*+/-*^ AGMs recovered slightly following explant, as there was now a statistically significant difference between heterozygous and homozygous knockout embryos; however, unlike with previously reported HSC numbers, progenitor numbers did not reach wild-type levels (**Figure 5D,E**). This was also the case with germline deleted embryos (**Figure S4C**,**D**). A substantial number of progenitors seem to remain in AGMs where both copies of *Gata3* were deleted from endothelial cells (**Figure 5B-E**); however, genotyping of individual colonies revealed that 6 out of 20 progenitors had escaped deletion (**Figure S4E**). This suggests that the effect on progenitors shown here is likely to be an underestimate, although germline-deleted embryos also retained some Gata3-independent progenitor activity (**Figure S4A-D**).

**Figure 5.**
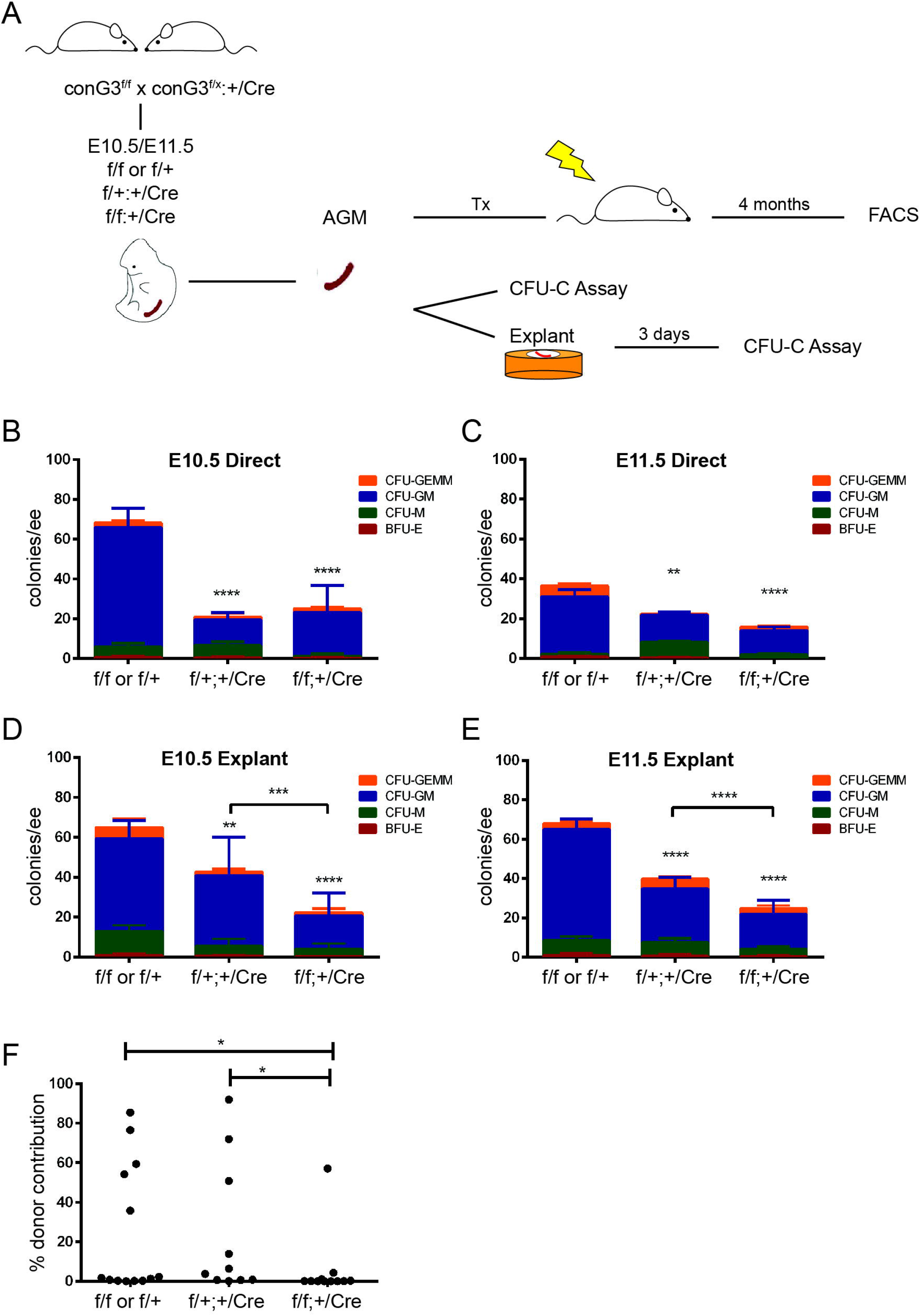
Gata3 is required for the endothelial-to-haematopoietic transition. (**A**) Outline of experiments performed with endothelial-specific (VEC-Cre) Gata3 knockout embryos. Colony numbers obtained with uncultured AGM cells from E10.5 (**B**) and E11.5 (**C**) embryos of the indicated genotypes. Colony numbers obtained with AGM cells from E10.5 (**D**) and E11.5 (**E**) embryos of the indicated genotypes following three days of explant culture. ** p<0.01; *** p<0.001; **** p<0.0001; Two-way ANOVA. (**F**) Percent donor contribution 4 months post-transplantation with E11.5 AGM cells of the indicated genotypes. * p<0.05; Mann-Whitney test.

AGMs were also transplanted to determine the effect of endothelial-specific *Gata3* deletion on HSC numbers. Repopulation activity was significantly reduced in homozygous knockout AGMs, demonstrating that Gata3 expression in the aortic endothelium is required for HSC generation (**Figure 5F**). In summary, our expression and functional data demonstrate an important cell-intrinsic role for Gata3 in regulating the transition of HECs to a haematopoietic fate.

## Discussion

This newly described cell-intrinsic role of Gata3 in the EHT adds another facet to the complex way in which Gata3 promotes HSC production in the AGM. We have previously demonstrated that Gata3 is expressed in two AGM niche compartments, the SNS ^19^ and the sub-aortic mesenchyme ^20^, from where it also supports emerging HSCs, albeit via different mechanisms. Arguably, the most essential function of Gata3 during development is the induction of catecholamine production in the SNS since the lack of this is the cause of mid-gestation lethality in Gata3-null embryos ^33^. This indirectly influences AGM haematopoiesis with secreted catecholamines supporting emerging HSCs ^19^. More recently, we have also described a role for Gata3 in the sub-aortic mesenchyme where it acts upstream of Runx1 ^20^. The relative contribution to AGM haematopoiesis of Gata3 in these three different compartments requires further dissection, although its role in HECs is likely to be the dominant one as the effect of the endothelial-specific knockout of Gata3 on HSPC numbers closely mirrors that of the germline knockout (^19^ and this manuscript). Yet, this defect was rescued through the administration of catecholamine derivatives ^19^, which could be explained by the fact that HSC production in Gata3-null AGMs is not entirely disrupted, with the remaining HSC activity being amplified through catecholamine addition.

Another interesting question is the relationship of Gata3 with the other two major regulators of HSC emergence, Gata2 and Runx1. We showed that Gata3 is required for Gata2 expression in the SNS, but not in the endothelium, and the expression of Runx1 in the AGM is reduced by half in Gata3-null embryos ^19, 20^; however, deletion of Runx1 and Gata2 from ECs has a much more profound negative impact on HSPC numbers ^7, 8^. Data from a recent scRNA-Seq analysis of aortic EC subpopulations shows an earlier upregulation of *Gata3* in arterial ECs as compared with *Runx1, Gata2* and *Gfi1* ^15^. Instead, it seems to closely mirror the expression pattern of *Hey2, Sox7* and *Sox17*, which need to be downregulated for HSCs to be able to mature ^34^. Why *Gata3* expression is restricted to a similarly brief window during EHT and whether sustained expression would also interfere with HSC formation remains to be investigated. Its co-expression with cell cycle inhibitors and correlation with changes in cell cycle is intriguing. *Gata3* has been shown to be expressed in adult quiescent LT-HSCs where it seems to regulate their entry into the cell cycle ^35, 36^. A recent scRNA-Seq dataset revealed that arterial EC and HEC are more quiescent than their venous counterparts and that HSPCs emerging at the end of the EHT start cycling again ^17^. This transient break in the cell cycle in HEC was reported in another gene expression study where it was also accompanied by a dip in metabolic activity ^16^. The subsequent increase in cycling as emerging HSC precursors mature is also supported by data from a cell cycle reporter mouse ^29^ and coincides with the stage at which *Gata3* becomes downregulated. scRNA-Seq analysis of human PSCs undergoing EHT also confirmed that ECs are largely quiescent, while HSPCs generated from these become more proliferative ^37^. Importantly, when cell cycle progression of ECs was chemically blocked, these cells could no longer generate HSPCs, implying that cell cycle entry is required to complete EHT. Taken all of these data into account, Gata3 may thus keep ECs/HECs quiescent while they undergo the major morphological changes required for the transdifferentiation into HCs and their release from the endothelial layer, and then, in analogy to its reported role in adult HSCs, may promote their re-entry into the cell cycle for completion of the EHT process.

## Acknowledgments

The authors are very grateful to the staff of the animal facilities both at the Cambridge Institute for Medical Research and the Centre for Regenerative Medicine for their support with animal experiments and to the flow cytometry teams at both of these institutes, Dr. Reiner Schulte, Dr. Chiara Cossetti, and Michal Maj in Cambridge and Fiona Rossi and Dr Claire Cryer in Edinburgh, as well as the Cambridge NIHR BRC Cell Phenotyping Hub, for excellent cell sorting services and help with flow cytometry analyses. We would further like to acknowledge the assistance of Matthew Gratian and Mark Bowen (Cambridge Institute for Medical Research) and Bertrand Vernay (Centre for Regenerative Medicine) with microscopy, and Drs Céline Souilhol and Stanislav Rybtsov with setting up the co-aggregate assays. We are also indebted to Jingfang Zhu and Meinrad Busslinger for providing the conditional Gata3 KO and the Gata3-GFP mice, respectively. Core facilities at the Edinburgh Centre for Regenerative Medicine were supported by centre grant MR/K017047/1. This work was funded by an Intermediate Fellowship from the Kay Kendall Leukaemia Fund (to K.O.), a Blood Cancer UK Bennett Senior Fellowship (10015 to K.O.) and a fellowship from the King Abdullah International Medical Research Centre (KAIMRC), Ministry of National Guard Health Affairs (to N.Z.). This research was also funded in part by the Wellcome Trust and the UKRI Medical Research Council. For the purpose of open access, the author has applied a CC BY public copyright licence to any Author Accepted Manuscript version arising from this submission.

## Author contribution

N.Z. performed and designed the majority of experiments; E.D. performed bioinformatics analyses; N.K.W. and B.G. provided advice and assistance with scRNA-Seq experiment; L.N. and A.F. performed experiments; L.M.F. provided important reagents; K.O. conceived and supervised the study and wrote the manuscript.

## Methods

### Mice

All animal work was carried out under a UK Home Office-approved licence and following local ethical approval. Males and females of Gata3-LacZ knock-in mice ^38^, Gata3-GFP knock-in mice ^21^, conditional Gata3 knockout mice ^32^, p57Kip2 knockout mice ^39^, VEC-Cre transgenic mice ^7^ and C57BL/6J mice were crossed to obtain embryos of the desired stage and genotype. The day of vaginal plug detection was considered as E0.5.

### AGM explant cultures

On the desired gestation day, the embryos were harvested in sterile medium (10% FCS (Hyclone) and 1% pen/strep in PBS (Sigma)). The developmental stage was specified by either counting somite pairs (E9.5-E10.5) or determining the stages by observing eye pigmentation (E11.5). Embryos smaller than their littermates or lacking a heartbeat were excluded. AGMs were dissected and cultured on Durapore filters (Millipore) at the air-liquid interface in M5300 long-term culture medium (Stem Cell Technologies) supplemented with 10^−6^ M hydrocortisone (Sigma). After 3 days, AGMs were dissociated with 0.125% collagenase (Alfa Aesar) in PBS/10% FCS at 37°C for 45minutes, followed by gentle pipetting, and single-cell suspensions were plated in methylcellulose.

### OP9 co-cultures

OP9 cells were maintained in αMEM (Gibco) with 20% heat deactivated FCS (Hyclone) and 0.22% sodium bicarbonate (Gibco) at 37°C with 5% CO_2_. 24h prior to the start of the co-cultures, cells were plated in αMEM with 10% FCS and 0.01% of 2-mercaptoethanol. Sorted haematopoietic and endothelial cell populations were plated on confluent OP9 stroma. Cultures were supplemented with SCF, FLt-3 ligand and IL7 at 10ng/ml concentration (all from Pepro-Tech) and incubated for 8–10 days at 37°C, 5% CO_2_. The haematopoietic nature of the cells generated in the OP9 co-cultures was assessed by CFU-C and flow cytometry.

### Co-aggregation cultures

AGM cells from E9.5-11.5 or E10.5 embryos were sorted and co-aggregated with OP9 cells according to the protocol by Rybtsov et al ^23^. Cell suspensions containing 1 embryo equivalent of the sorted cells and 10^5^ OP9 cells in 30μl volume of media (Iscove’s modified Dulbecco’s medium [IMDM], Invitrogen-GIBCO, 20% of heat-inactivated FCS, L-glutamine, penicillin/streptomycin) were centrifuged at 450 × g for 12min in 200μl pipette tips sealed with parafilm. The IMDM medium for the culture steps was supplemented with SCF, Flt3 and IL3 at a concentration of 100ng/ml (all from Pepro-Tech). Co-aggregates were cultured on floating 0.8mm AAWP 25 mm nitrocellulose membranes (Millipore) for 4–5 days at 37°C. Cultured aggregates were dissociated using collagenase and either analysed by flow cytometry or transplanted into irradiated mouse recipients.

### Colony-forming assays

Dissociated AGM cells (either fresh or after explant culture) were added to 3.6ml methylcellulose (M3434; Stem cell technologies) and plated in triplicates (65.000 cells/plate). Plates were incubated at 37°C and haematopoietic colonies counted and scored 7 days later.

To detect *Gata3* deletion by VEC-Cre, individual colonies were picked from methylcellulose plates and added to 50μl of Alkaline lysis buffer (0.04% disodium EDTA and 0.25% NaOH in water), followed by 20min incubation at 95°C, 280 × g shaking. 50 μl of Neutralisation Reagent (4% 1M Tris-HCl in water) was then added to terminate the reaction. 10μl of this was added to 25μl PCR mastermix, comprised of 12.5μl KAPA2G Fast ReadyMix with dye (Anachem) and 2.5μl of primer mix, using the following primers: forward, CAGTCTCTGGTATTGATCTGCTTCTT; and reverse, GTGCAGCAGAGCAGGAAACTCTCAC. PCR conditions were 5min at 94°C, 30 cycles each of 20sec at 94°C, 20sec at 55°C and 20sec at 72°C, followed by a final incubation of 5min at 72°C.

### Transplantations

Single-cell suspensions were resuspended in 500μl PBS/2% FCS and injected intravenously into irradiated recipients together with 2 × 10^5^ spleen cells (for direct transplantations) or 2 × 10^4^ bone marrow cells (for co-aggregates) to alleviate the irradiation effects. Recipients were irradiated with a split dose of 460-475 rad with three hours between each dose, using a Caesium source. The congenic CD45 system was used with allelic variants of CD45 (CD45.1 and CD45.2) to distinguish donor from recipient haematopoietic cells. Recipients were CD45.1/.2 or CD45.1/.1 and the donors were CD45.2/.2 on a C57BL6J background. Recipient blood was analysed at 1 and 4 months post-transplantation and considered positive for reconstitution if the donor contribution was more than 5%.

### Flow cytometry

Antibody stainings were performed in FACS buffer (2% FCS in PBS) for 30min on ice in the dark, using the following antibodies: anti-CD41-BV421 (1:100; BD Bioscience), anti-CD45-BV421 (1:100; BD Bioscience), anti-CD45-APC-Cy7 (1:100; BD Bioscience), anti-CD45-A700 (1:50; Biolegend), anti-CD45.1-PE (1:200; Biolegend), anti-CD45.2-A700 (1:200; Biolegend), anti-CD43-BV421 (1:100; BD Bioscience), anti-Ter119-V500 (1:100; BD Bioscience), anti-VEC-PE-Cy7 (1:100; Biolegend), anti-VEC-AF647 (1:100; BD Bioscience), anti-CD11b-PB (1:200; Biolegend) and anti-Gr1-PB (1:200; Biolegend). Peripheral blood samples from transplant recipients were pre-treated with Red Cell Lysis buffer (BD Bioscience). Dead cells were excluded either via 7-aminoactinomycin D staining 1:1000 (7AAD, Invitrogen) or Sytox AAD (1:5000) (Invitrogen). All experiments included the following controls: unstained cells, single stained sample for each flurochrome and fluorescent minus one (FMO) controls. Cells were analysed using LSRFortessa (BD Bioscience) or sorted using MoFlo (Beckman Coulter), ARIA (BD Bioscience), or Fusion (BD Bioscience) and data analysed with the FlowJo software (BD Bioscience).

For cell cycle analysis, sorted cells were incubated for 1min at room temperature 1:1 with DAPI staining solution (5ug/ml DAPI (Sigma) and 1% (v/v) Nonidet P40 (Sigma) in dH2O) and analysed on an LSRFortessa (BD Bioscience).

### Immunohistochemistry

Embryos were fixed in 2% paraformaldehyde (Sigma) in PBS for 1.5 hour at 4°C, while rotating, then washed three times in PBS, and cryoprotected overnight in 30% sucrose/PBS at 4°C, rotating. Fixed embryos were embedded in OCT TissueTek Compound, quick-frozen on dry ice and stored at -80°C. 10μm sections were prepared on a cryostat (Leica, CM3050 S).

For antibody staining, either an Avidin/biotin system or fluorescent-labelled secondary antibodies were used. Cryosections were washed three times with PBS/1% FCS/0.1% Triton-X100 for five minutes each and blocked with 200μl of PBS/0.05% Tween/1% BSA, containing Avidin/biotin block where appropriate. Slides were incubated with the primary antibody in PBS/0.05% Tween/1% BSA for 24 hours at 4°C in the dark. After the incubation, the slides were washed with PBS/1% FCS/0.1% Triton-X100, before being incubated with the secondary antibody or fluorescently labelled streptavidin in PBS/0.05% Tween/1% BSA for 45 minutes at room temperature in the dark. The slides were washed with PBS/1% FCS/0.1% Triton-X100 and mounted with Vectashield containing DAPI (Vectorlabs). The following antibodies were used: anti-Th (mouse; 1:300; Millipore), anti-GFP (chicken; 1:400; Thermofisher), anti-GFP (rabbit; 1:500; Life Technologies), FITC-labelled anti-CD34 (rat; BD Bioscience; 1:100), anti-Gata3 (goat; 1:300; BD Bioscience), Alexa647-labelled anti-chicken (1:500; Jackson Immunoresearch), Alexa546-labelled anti-mouse (1:200; Life Technologies), Alexa647-labelled anti-rabbit (1:200; Life Technologies), Alexa555-labelled anti-rabbit (1:500; Invitrogen), and Alexa633-labelled anti-goat (1:200; Sigma). Images were acquired on a Leica SP8 confocal microscope and analysed with Leica Las × software.

### X-gal staining

Gata3^lz/+^ embryos were washed with PBS/0.02% NP40 and fixed for 1 hour with PBS/10% Formal Saline/0.2% Glutaraldehyde/2mM MgCl2/5mM EGTA/0.02% NP40 at 4°C. They were then washed three times for 30 minutes each with PBS/0.02% NP40 and stained overnight at room temperature with 5mM K3Fe(CN)6/5mM K4Fe(CN)6/2mM MgCl2/0.01% Nadeoxycholate/0.02% NP40/0.1% X-gal (all in PBS). Stained embryos were cryopreserved and sectioned as above and sections counterstained with Neutral Red.

### Endogenous gene activation

#### Human iPSCs maintenance

Human iPSCs were cultured in StemPro hESC SFM (Gibco) supplemented with 20ng/ml bFGF (R&D) on CELLstart (Gibco) coated wells. Media was changed daily and the cells passaged every 3–4 days at a ratio of 1:4 using the StemPro EZPassage tool (ThermoFisher Scientific).

#### Human iPSC transfection

Single cell suspension was obtained using Accutase (Gibco) and 3⍰×⍰10^5^ cells were reverse transfected with 2μg of UniSAM DNA using the Xfect Transfection reagent (Clontech) and plated into a coated 6 well plate. Each well was transfected with the PB-UniSAM plasmid (Addgene 99866 {Fidanza, 2017 #17}) containing either one of the four gRNAs against RUNX1C promoter {Fidanza, 2017 #17} or no guide (Empty vector control).

#### Gene expression analysis

The total RNA was extracted using the RNAeasy Mini Kit (Qiagen) 2 days post transfection and the cDNA was synthesised from 500ng of total RNA using the High-Capacity cDNA synthesis Kit (Applied Biosystem). 2ng of cDNA were amplified per reaction and each reaction was performed in triplicate using the LightCycler 384 (Roche) with SYBR Green Master Mix II (Roche), β-Actin was used as reference genes. Gene activation values were calculated as fold change relative to the empty vector control group. Targets and reference genes were amplified using the following primer pairs: RUNX1C_fw agcctggcagtgtcagaagt, RUNX1C_rv gggactcaatgatttcttttacca, GATA3_fw gctcttcgctacccaggtg, GATA3_rv gtaaaaaggggcgacgactc, ACTB_fw ccaaccgcgagaagatga, ACTB_rv ccagaggcgtacagggatag.

### RNA sequencing

#### Cell sorting

Libraries for RNA sequencing were prepared according to the protocol by Picelli et al. ^40^. Cells were initially sorted from 3-5 biological replicates into tubes based on their populations: Gata3-GFP+ EC, Gata3-GFP-EC, Gata3-GFP+ HC and Gata3-GFP-HC. Each population was then sorted again into 96 wells plate, 20 cells/well (20 pools per population), containing 2.3μl of lysis buffer (0.2% RNase inhibitor (Ambion, Thermo Fisher Scientific) in Triton X-100 (Sigma)). Plates were spun down and stored at -80°C until further processing.

#### Reverse transcription

For the reverse transcription step, 2μl of annealing mix (5% ERCC RNA spike-In Mix (pre-diluted at 1:25,000; Invitrogen), 5% Oligo-dT (5⍰–AAGCAGTGGTATCAACGCAGAGTACT30VN-3⍰; 100μM; biomers.net), 50% dNTP 10mM (Fermentas) and 40% distilled water) were added to each well and the plates incubated at 72°C for 3 minutes and immediately placed on ice. 5.7μl of reverse transcription mixture (0.5μl Superscript II RT (200 U/μl; Invitrogen), 0.25μl RNase inhibitor (20 U/μl), 2μl 5x Superscript II First Strand Buffer (Invitrogen), 0.5μl DTT (Invitrogen), 2μl 100μM Betaine (Sigma), 0.06μl 1M MgCl2 (Ambion), 0.1μl TSO Oligo (5⍰-AAGCAGTGGTATCAACGCAGAGTACATrGrG+G-3⍰; 100 μM; Exiqon) and 0.29μl distilled water) were then added per well and the plate placed in a PCR cycler with the following settings: 42°C for 90 minutes, 10 cycles of: 50°C for 2 minutes, 42°C for 2 minutes, then at the end 70°C for 15 minutes.

#### PCR Pre-amplification

For the PCR Amplification, 15μl of the PCR mixture were added per well, consisting of 12.5μl KAPA HiFi Hotstart ReadyMix (2x; KAPA Biosystems), 0.25μl ISPCR primer (5⍰-AAGCAGTGGTATCAACGCAGAGT-3⍰; 10 μM; biomers.net) and 2.25μl distilled water. The reactions were run at 98°C for 3 minutes, 21 cycles of: 98°C for 20 seconds, 67°C for 15 seconds, 72°C for 6 minutes, and 72°C for 5 minutes at the end.

Ampure XP beads (Beckman Coulter) for PCR clean up were added to each sample at a 1:1 ratio mixture and the mixture incubated at room temperature for 8 minutes before fitting the plate on a plate magnetic stand. The plate was left on the stand for 5 minutes until all beads had been collected at one well corner. The supernatant was then discarded, and the beads washed twice with 200μl of 80% ethanol for 30 seconds each time. The samples were recovered with 20μl of Elution Buffer (Qiagen). The size distribution of the cDNA library was checked on an Agilent high-sensitivity DNA chip (Agilent Technologies), according to the manufacturer instructions.

#### Sequencing library preparation

Tagmentation was carried out using the Illumina Nextera XT DNA sample preparation kit (Illumina) according to an optimised Tagmentation protocol (Fluidigm). 1.25µl of the eluted cDNA was mixed with 2.5µl tagmentation DNA buffer and 1.25µl Amplification Tagment mix and the reactions placed in a thermal cycler for 10 minutes at 55°C. NT buffer was then added at 20% ratio to the original volume in each well and the plate centrifuged at 4,000 rpm for 5 minutes. Nextera PCR Master Mix (NMP) was added at a ratio of 37.5% to the initial volume, and Index Primers 1 (N701-N712) and 2 (S501-S508) at a ratio of 12.5% (each) were combined in a way that each well was uniquely labelled and dual-indexing metadata could be obtained (Nextera XT 96-Index kit; Illumina). PCR amplification was performed in a thermal cycler at these settings: 72°C for 3 minutes, 95°C for 30 seconds, 12 cycles of: 95°C for 10 seconds, 55°C for 30 seconds, 72°C for 60 seconds and 72°C for 5 minutes. Libraries were pooled with 1μl from each well and cleaned up by mixing with Ampure XP beads at 70% of the total pool volume. After two washes with 1.2μl of 80% ethanol, libraries were eluted in 50μl of elution buffer. The library size distribution was checked on an Agilent high-sensitivity DNA chip and the library quantified using the KAPA library quantification kit (KAPA Biosystems). For each reaction, 6μl of ‘2X KAPA SYBR FAST qPCR Master Mix +10X Primer Premix’ was mixed with 2μl PCR-grade water and 2μl of diluted library. Each library was diluted in DNA Dilution Buffer (10mM Tris-HCl, pH 8.0 and 0.05% Tween-20) at 1:2,000,000 and run in triplicates along with the provided DNA Standards (Standards 3-6 were used) and No Template Controls. The following settings were used: 95°C for 5 minutes, 35 cycles of: 95°C for 30 seconds, 60°C for 45 seconds and 65-95°C for a melting curve step. For the data analysis, a standard curve was generated and the concentration was calculated for each library. Pooled libraries were sequenced on an Illumina Hi-Seq 4000 (Sanger, Cambridge), as single-end 125 base pair reads.

#### Sequencing data analysis

The data were aligned using STAR ^41^ to Ensembl genome build 81 ^42^, with gene counts obtained using HT-Seq ^43^. Quality control filtering and normalisation was performed in R. More than 500,000 reads uniquely mapped (either to ERCC spike-ins or endogenous mRNA), with more than 20% of total reads mapped to mRNA, less than 20% of mapped reads allocated to mitochondrial genes, less than 20% of reads mapped to ERCC spike-ins and more than 8000 high coverage genes. Cells were normalised with Scran ^44^ and highly variable genes identified estimating technical variance with the ERCC spike-ins to ^45^. In-house programs in R were used for PCA and t-distributed Stochastic Neighbor Embedding (t-SNE) dimensionality reduction. Genes differentially expressed between cell types and Gata3 expression groups was performed using the rank_genes_groups function with the t-test_overestim_var method. P-values were adjusted using the benjamini-hochberg procedure, and genes with adjusted p-value < 0.01 considered as significant.

Gene Ontology (GO) analysis was performed using the Gene Ontology Consortium Enrichment Analysis, which utilizes PANTHER Classification System for biological processes in mus musculus. The data have been deposited in NCBI’s Gene Expression Omnibus (GEO) repository under accession number GSE114926.

### Statistical Analysis

Graph preparations and statistical analysis were performed using GraphPad Prism. The Mann-Whitney test was used to determine significance levels for transplantation experiments, paired t-test was used to determine the significance levels for colony forming assay following OP9 co-culture and co-aggregates, and two-way ANOVA test was used to determine significance levels for colony-forming assay.

## Supplementary Figures

**Figure S1 Gata3-GFP expression mirrors that of endogenous Gata3 protein**.

Cryosection from a G3-GFP+ E11.5 embryo showing the co-staining between Gata3 (green) and GFP (red) with DAPI as nuclear stain in blue. (**A**) merged image, (**B**) antibody to GFP, (**C**) antibody to Gata3. (**D-F**) Magnification of area indicated by yellow box in B and C demonstrating co-staining between Gata3 and GFP. DA: dorsal aorta, AG: adrenal anlage, SG: sympathetic ganglia.

**Figure S2 Gata3-GFP+/-EC and HC sorting strategy**.

(**A**) Gating strategy for flow cytometry sorting of Gata3-GFP+/-ECs (VEC+ CD41/45-) and HCs (CD41/45+) for the co-culture experiments. (**B**) Gating strategy for flow cytometry sorting of Gata3-GFP+/-ECs (VEC+ CD41/43/45-Ter119-) and HCs (CD41/43/45+ Ter119-) for the co-aggregate experiments.

**Figure S3 Gene ontology terms enriched in differentially expressed genes**.

GO terms significantly enriched amongst the genes upregulated and downregulated in Gata3-GFP+ ECs and HCs

**Figure S4 Colony-forming assays with Gata3 germline-deleted AGMs**.

CFU-A assays of *Gata3*^*+/+*^ (WT), *Gata3*^*+/-*^ (HET) and *Gata3*^*-/-*^ (KO) uncultured E10.5 (**A**) and E11.5 (**B**) AGM cells and following 3 days of explant culture of E10.5 (**C**) and E11.5 (**D**) AGMs. ** p<0.01; *** p<0.001; **** p<0.0001; Two-way ANOVA. (**E**) Gel electrophoresis result of colonies demonstrating that some cells retained the floxed allele and therefore had escaped recombination and deletion of *Gata3*.

