## Supplementary figures and images for "Endothelial-specific Gata3 expression is required for haematopoietic stem cell generation"

### Supplemental Figure 1

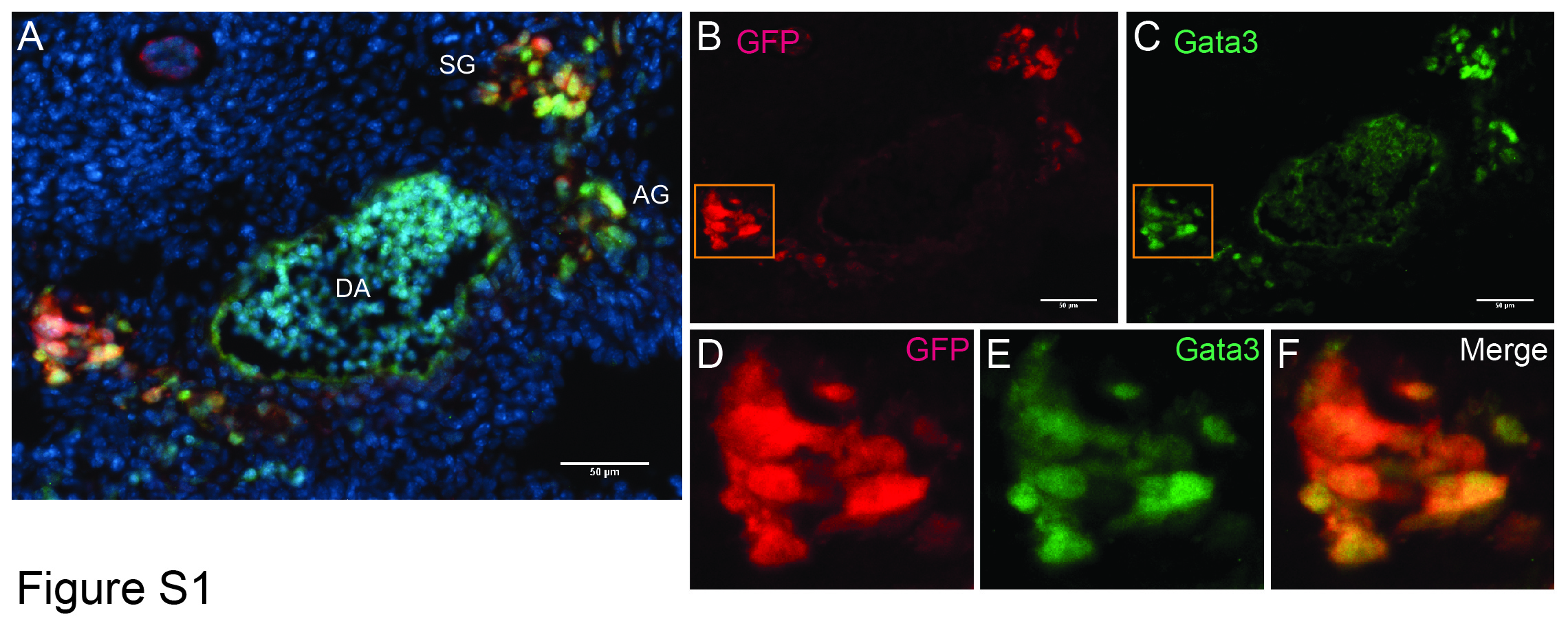

### Supplemental Figure 2

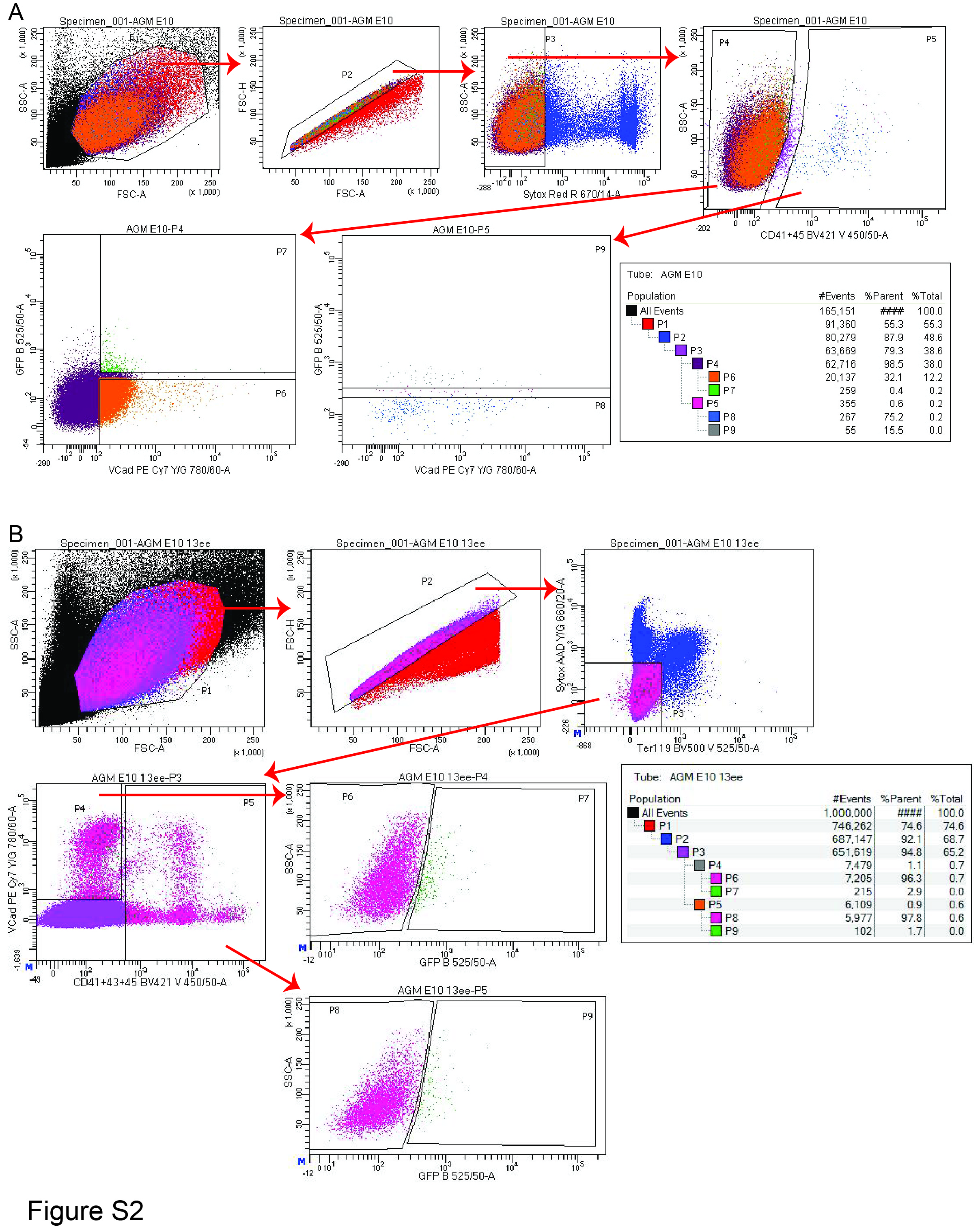

### Supplemental Figure 3

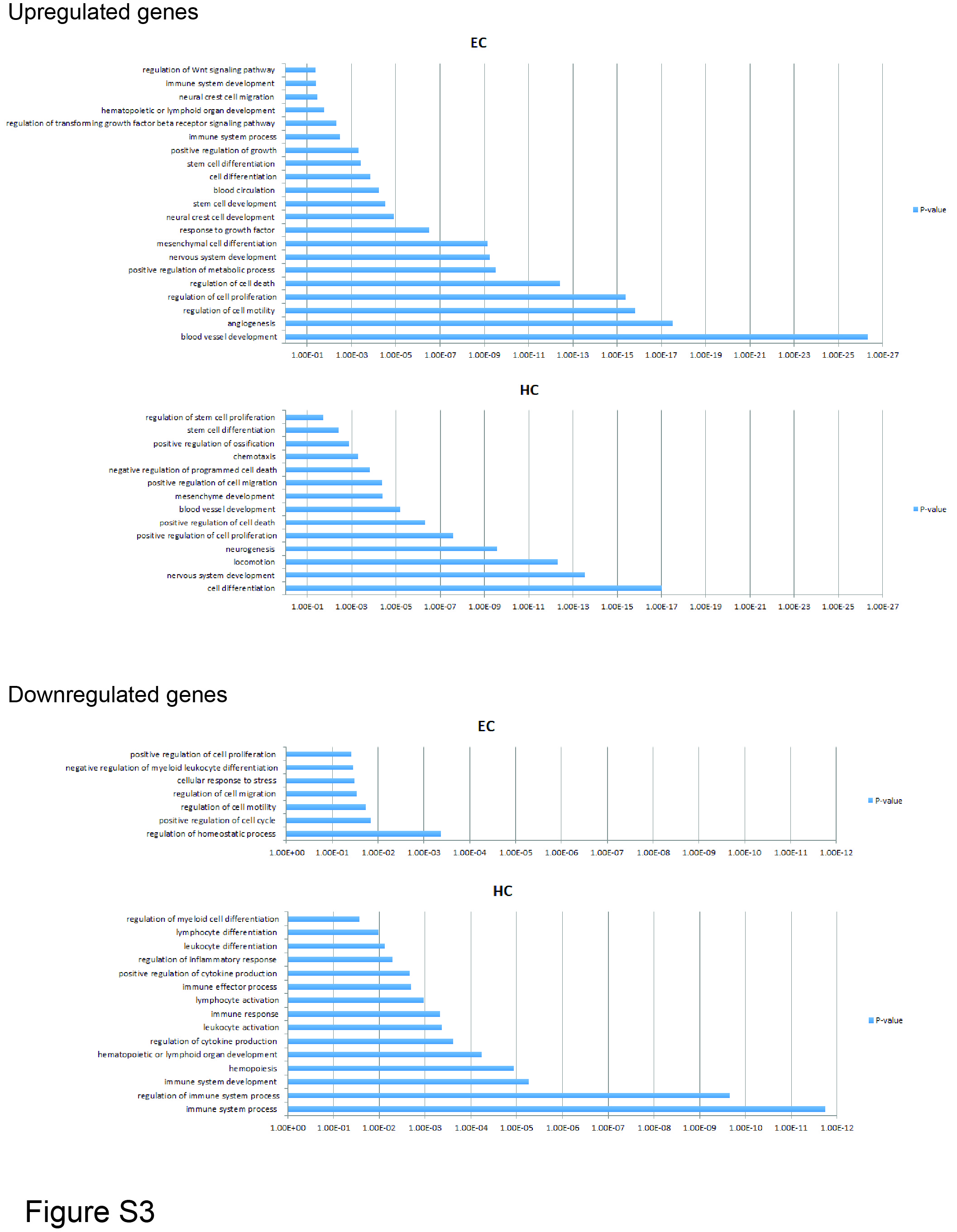

### Supplemental Figure 4

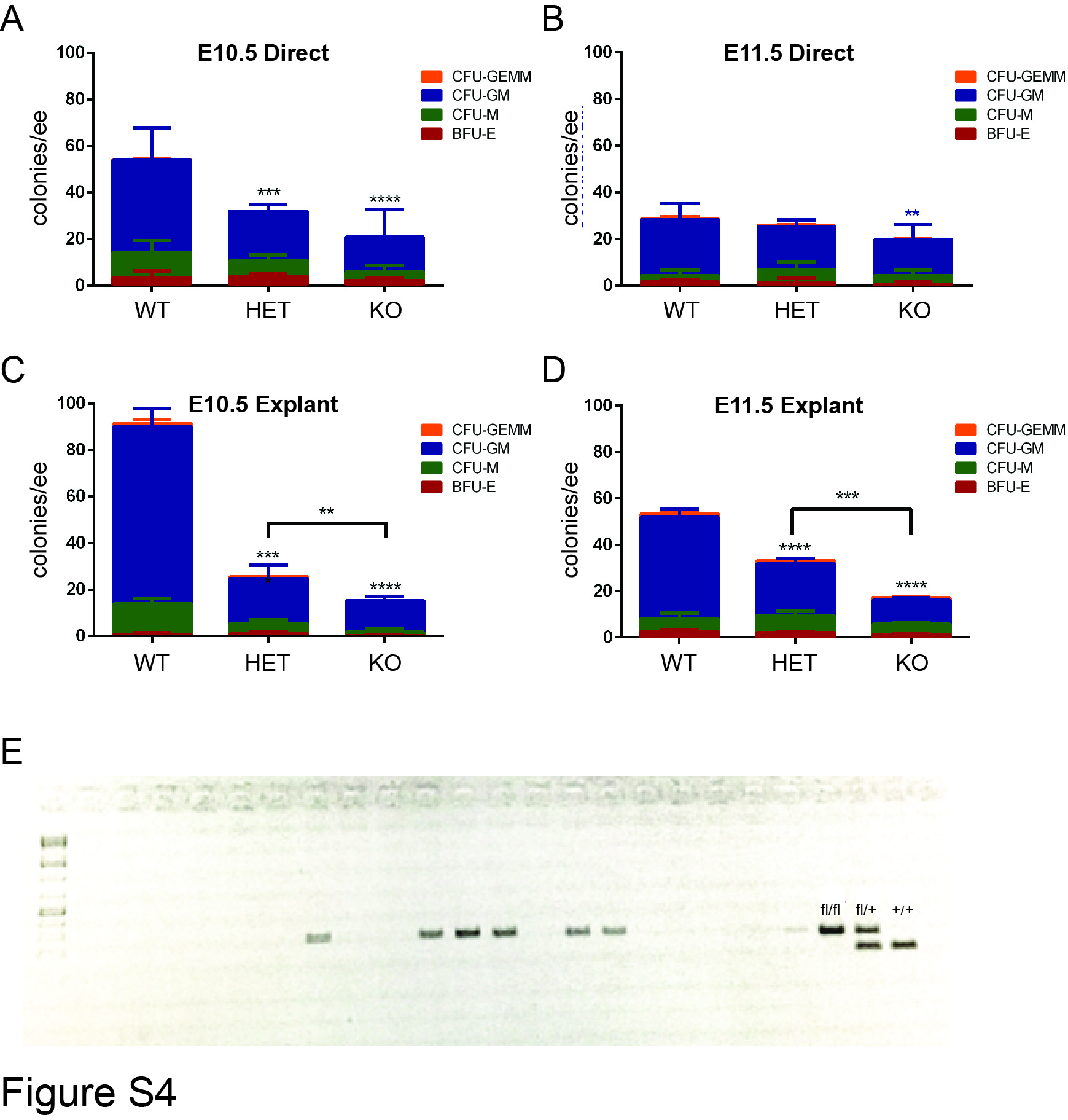
